# Stochastic Single Cell Behaviour Leads to Robust Horizontal Cell Layer Formation in the Vertebrate Retina

**DOI:** 10.1101/561449

**Authors:** Rana Amini, Anastasia Labudina, Caren Norden

## Abstract

Developmental programs that arrange cells and tissues into patterned organs are remarkably robust. In the developing vertebrate retina for example, neurons reproducibly assemble into distinct layers giving the mature organ its overall structured appearance. This stereotypic neuronal arrangement, termed lamination, is important for efficient neuronal connectivity. While retinal lamination is conserved in many vertebrates including humans, how it emerges from single cell behaviour is not fully understood. To shed light on this question, we here investigated the formation of the retinal horizontal cell layer. Using *in vivo* light sheet imaging of the developing zebrafish retina, we generated a comprehensive quantitative analysis of horizontal single cell behaviour from birth to final positioning. Interestingly, we find that all parameters analyzed including cell cycle dynamics, migration paths and kinetics as well as sister cell dispersal are very heterogeneous. Thus, horizontal cells show individual non-stereotypic behaviour before final positioning. Yet, these initially stochastic cell dynamics always generate the correct laminar pattern. Consequently, our data shows that lamination of the vertebrate retina contains a yet underexplored extent of single cell stochasticity.

## INTRODUCTION

During development, tissues and organs change their morphology to acquire their characteristic sizes, shapes and cellular patterns that ensure correct function. These processes are tightly regulated to yield reproducible outcomes and precise tissue patterning. The vertebrate retina is a fascinating example of such pattern formation. As an extension of the central nervous system, it develops from a pool of multipotent progenitors into a neural tissue composed of five neuronal cell types. Neurons of the same type as well as the one glial cell type, Müller glia, are positioned in distinct layers giving the mature retina its laminated appearance (Figure 1A). Interestingly, retinal development contains stochastic elements. For example, stochasticity in the apical–basal movement of neuroepithelial cell nuclei was proposed to contribute to proliferative versus neurogenic division outcome in zebrafish (Baye and Link, 2007; Del Bene et al., 2008). In addition, stochasticity was suggested to be involved in retinal neuronal fate determination (He et al., 2012). While studies so far concentrated on exploring stochasticity of fate decisions, whether other phenomena, particularly retinal lamination, also show stochastic elements remained elusive. To answer this question, we explored single cell behaviour of horizontal cells (HC) before lamination. HCs are inhibitory interneurons that localize beneath the photoreceptor cell (PR) layer and form synaptic connections with PRs and bipolar cells (BCs) (Amini 2018). BCs ultimately transmit visual information to the retinal ganglion cells (RGCs) (Figure 1A). Committed HC precursors (HCpr) divide once more before integrating into their laminar array producing two HCs (Godinho et al., 2007; Weber et al., 2014). The timing and dynamics of this last cell cycle have not yet been studied. Furthermore, HC migration is not fully understood. Unlike RGCs that keep an apical connection and reach their final destination via stereotypical somal translocation (Icha et al., 2016), HCs are not attached but instead undergo multipolar bidirectional migration to reach their final position (Chow et al., 2015; Edqvist and Hallböök, 2004). While different phases of movement have been characterized (Chow et al., 2015), the stereotypicity of these movements remained unexplored.

**Figure 1:**
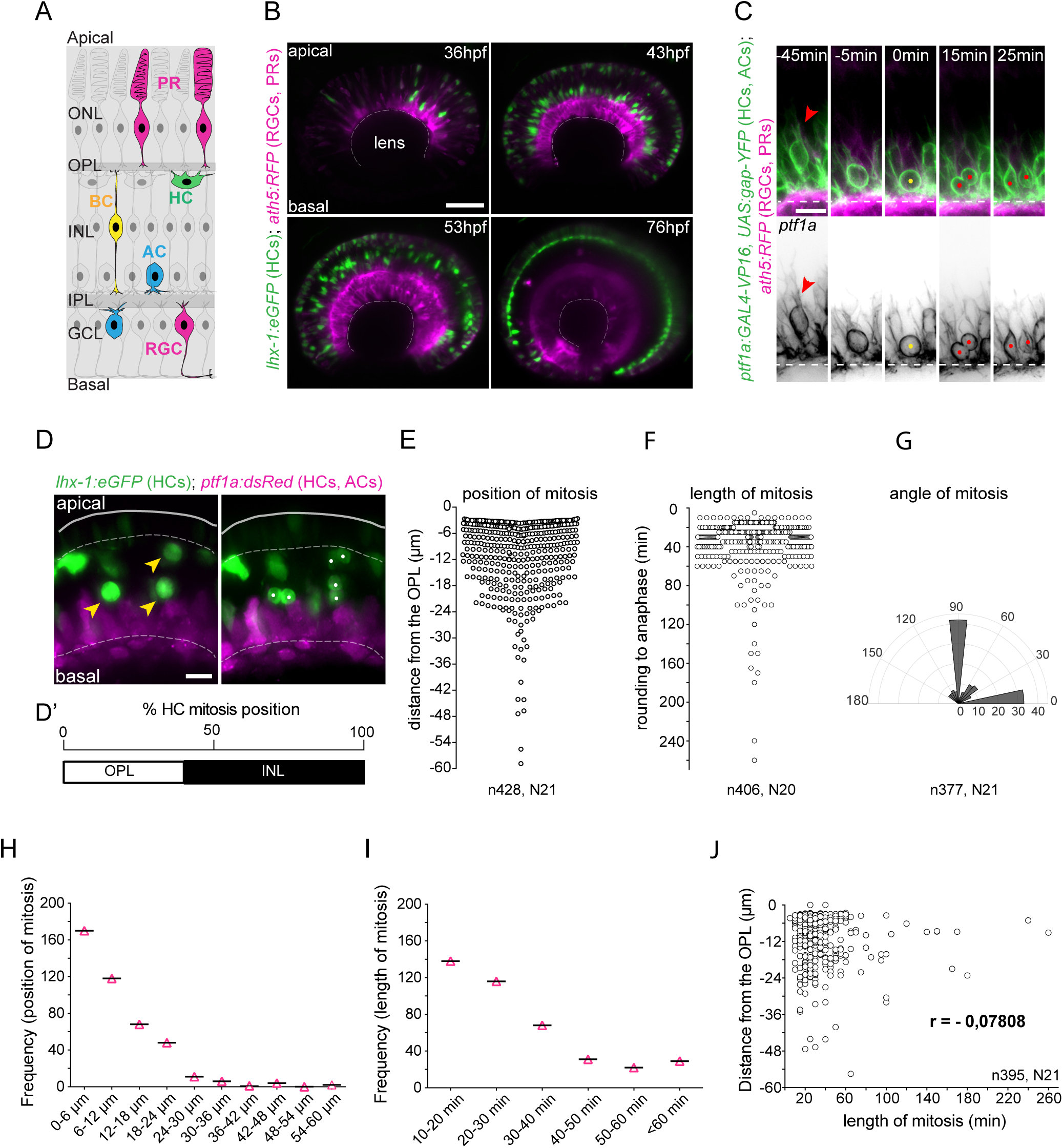
Heterogeneity in mitotic position and behaviour of HCprs. **(A)** Scheme of zebrafish retina. Neurons: photoreceptors (PR, magenta); horizontal cells (HC, green); bipolar cells (BC, yellow); amacrine cells (AC, blue) and retinal ganglion cells (RGC, magenta). Layers: outer nuclear layer (ONL), inner nuclear layer (INL) and ganglion cell layer (GCL), outer plexiform layer (OPL) inner plexiform layer (IPL). **(B)** Montage of HC lamination 36hpf-76hpf. *lhx-1:eGFP* (HC and PR, green), *ptf1a:dsRed (AC and HC, magenta)*, Scale Bar, 10 µm. (**C)** Montage representing a typical example of HCpr mitosis length. Time is relative to mitosis onset. Red arrowhead: pre-mitotic protrusive cell, yellow dot: mitotic cell, red dots: sister cells after division. Dotted white line: IPL. Scale Bar, 5 µm. **(D)** Position of HCpr mitosis is heterogenous. Yellow arrowheads: mitotic HCprs, white dots: sister HCs after division. White dotted line: OPL (top), IPL (bottom). Scale Bar, 10 µm. **(D’)** Percentage distribution of HCpr divisions. **(E)** position (µm), **(F)** length (min) and **(G)** angle of HCpr mitosis. Frequency of mitotic **(H)** position and **(I)** length. **(J)** Plot of mitotic length versus position shows no strong correlation. Regression analysis using Pearson correlation (r).

We here used long-term *in vivo* light sheet imaging and quantified single cell behaviour over a large cohort of HCs following cell-cycle parameters and bidirectional migration (Chow et al., 2015; Godinho et al., 2007; Weber et al., 2014). Surprisingly, we find that all parameters substantially vary between single cells before robust final lamination. Thus, we provide the first evidence that, as seen for fate decisions, also retinal lamination contains stochastic elements.

## RESULTS

### HCprs vary in mitotic position, length and spindle orientation

Before final lamination, committed HCprs undergo one more division. The position of this division is not restricted to the final HC location but can also occur along the inner nuclear layer (INL) (Godinho et al., 2007; Weber et al., 2014). While previous studies used immunostaining of fixed samples, no appreciation of HCpr division dynamics existed.

Following 428 cells from 21 embryos (from here on n=428, N=21) (as experiment example see Movie 1 and Figure 1B), we found that, as postulated previously (Godinho et al., 2007; Weber et al., 2014), all HCprs underwent an additional division resulting in two HCs that both later entered the HC layer (data from all pooled movies, S3 Table). None of the tracked HCprs underwent more than one terminal division (data from all pooled movies S3 Table). Single cell analysis of mitotic location (Figure 1D, D’) (n=428, N=21) revealed variations in HCpr mitotic position. While most cells entered and underwent mitosis near the outer plexiform layer (OPL) where they later resided, divisions were also seen along the whole apico-basal length of the INL, even close to the inner plexiform layer (IPL) (Figure 1C, 1D, 1D’ and 1E). All markers (see Material and Methods) showed a similar mitotic position pattern (Supplementary Figure 1E). Only in very rare cases did HCprs divide in the RGC layer (Supplementary Figure 2A). Frequency analysis confirmed a bias towards OPL positions as 41% of HCpr divisions occurred close to the OPL (distance = 0 µm – 6 µm), whereas the remaining 59% spread along the INL (Figure 1D’, 1E and 1H). This suggests that HCprs favor divisions closer to the prospective HC layer. While this bias was not shown previously, it was postulated that divisions close to the OPL facilitate rapid synaptic contacts between HCs and their partners (Godinho et al., 2007).

To understand whether these positional differences were connected to differences in HCpr mitotic behaviour, we analyzed the duration of mitosis from rounding to anaphase onset (Figure 1C and 1F). Single cell and frequency distribution analysis (n=404, N=20) revealed that while 62% of HCprs divided within the first 30 min after rounding, mitosis length showed a wide variety from 10 min to 260 min (Figure 1F and 1I). This is markedly different from what was reported for retinal progenitor cells that have very reproducible mitotic timing lasting 25 +/- 3min (Leung et al., 2011). To reveal potential differences between cells with short versus long mitotic length, we used a nuclear marker to differentiate between chromatin condensation, metaphase plate formation and DNA segregation. We found that the time between metaphase and anaphase was similarly diverse as overall mitotic length (Supplementary Figure 2B). Cells that took longer between metaphase plate formation and final division showed intense spindle rocking before anaphase onset (Supplementary Figure 2C, Movie 2). Moreover, final division angles had a wide distribution variety (n=377, N=21) (Figure 1G), suggesting that HCprs can divide with any directionality. Nevertheless, a bias towards 0° and 90° division angles was seen. No strong correlations were observed between division angle and timing of spindle positioning (Supplementary Figure 2D and E). In addition, the position of mitotic cells showed no strong connection to mitotic length (Figure 1J), suggesting that duration of spindle positioning was the main factor in mitotic length distribution.

Together, our data shows that division position, duration of final mitosis and spindle orientation are not stereotypic but vary widely among HCprs. However, independently of these variations resulting HCs robustly reach their correct layer (Figure 1B, Movie 1).

### HCprs exhibit heterogenous cell cycle timing

The fact that divisions of HCprs can in principle occur at any position within the INL made us ask whether these cells follow an intrinsic cell cycle timer making cell divisions occur independently of HCpr position. To test this idea, we tracked HCprs from apical birth to final division (n=23, N=10) (Figure 2A, Movie 3). Cell cycle duration ranged from 11h15min to 16h41min, with an average of 13h (Figure 2C). Thereby, HCpr cycles were much longer than what was reported for progenitor cells that, albeit showing variations, have an average cell cycle length of 7.3h at 30 hpf and 5.3h at 42 hpf (Matejčić et al., 2018). As we used light sheet imaging to analyze cell cycle timing, it is unlikely that differences were due to increased phototoxicity (Icha et al., 2016; Icha et al., 2017). No clear correlation between cell cycle length and distance from apical surface during division was seen (Figure 2D).

**Figure 2:**
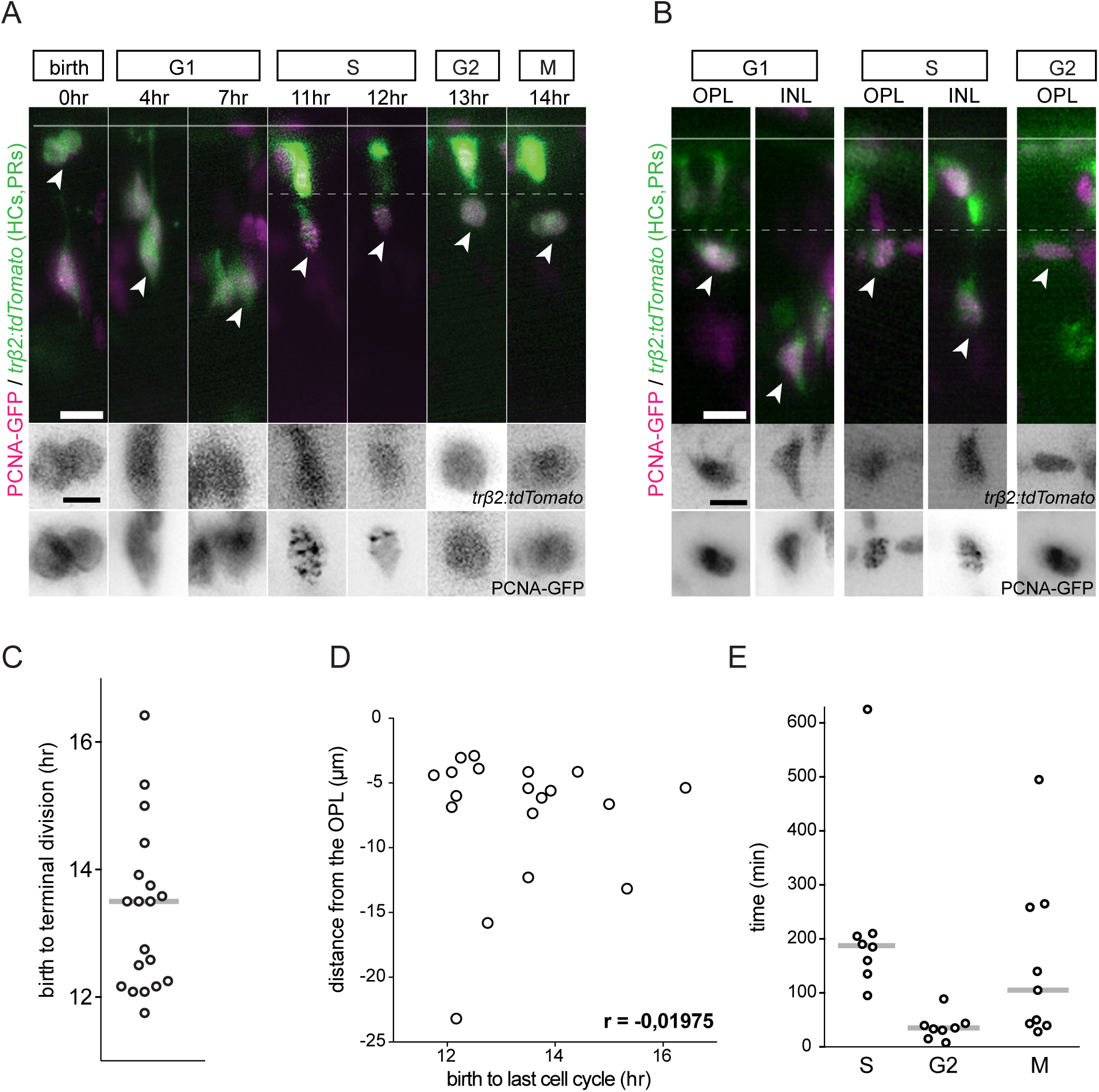
HCprs undergo long heterogenous cell cycles. **(A)** Montage of HCpr cell cycle. *Trβ2:tdTomato* (HCpr, green) and PCNA-GFP (nuclei, magenta). Upper Panel; white arrowheads: tracked cell. Dotted white line: OPL. Scale Bar, 10 µm. Inlays in bottom panels: *Trβ2:tdTomato* and PCNA-GFP, respectively. Scale Bar, 5 µm. **(B)** HCpr cell cycle phases occur at different positions in INL. *Trβ2:tdTomato* (HCpr, green), PCNA-GFP (nuclei, magenta), white arrowheads: cell followed. Top panel: scale bar, 10 µm. Inlays in bottom panels, *Trβ2:tdTomato* and PCNA-GFP, respectively. Dashed white line: OPL. Scale Bar, 5 µm. **(C)** Distribution of HCpr cell cycle length. Line: mean. **(D)** Scatter plot of HCpr cell cycle length versus HCpr position shows a lack of strong dependence. Regression analysis using Pearson correlation (r). **(E)** Distribution of HCpr cell cycle phase length, S, G2 and M. Line: mean.

Studies of fixed chick retinae suggested that HCprs undergo S phase during apical-basal migration and arrest in G2 while stalling in the INL (Boije et al., 2009). We tested this notion using PCNA (Proliferating cell nuclear antigen) as a marker to unambiguously follow HCpr cell cycle phases (Leung et al., 2011) (Figure 2A and 2B and Movie 3). Interestingly, we found that HCprs passed G1 and S phase at diverse positions along the INL (Figure 2B). G1 can occur in the INL when cells are already multipolar and S phase occurred during basal as well as apical migration (Figure 2B). In addition, we found that G2 (defined as time between disappearance of PCNA dots and cell rounding (Leung et al., 2011)) covered a rather short span of the overall cell cycle length (Figure 2E) often occurring close to the OPL (Figure 2B).

Together, this data makes it unlikely that HCprs arrest in G2 before apical migration in zebrafish. Further, our results argue against an intrinsic cell cycle timer responsible for mitotic onset (Figure 2D).

### HC migration is non-stereotypic and varies in length and overall trajectories

It was previously reported that maturing HCs undergo bidirectional migration before final lamination (Chow et al., 2015; Edqvist and Hallböök, 2004; Weber et al., 2014). However, so far only small cohorts of cells were analyzed using confocal microscopy with time resolution of 15 minutes or longer and the focusing on different movement phases (Chow et al., 2015). To understand the stereotypicity of HC single cell migration, we followed them from birth to final positioning using light sheet microscopy at 5min intervals (from here on we will use the term HC for all migration behaviour before and after final division) (n=21 cells N=11) (Figure 3A, Supplementary Figure 3A, Movie 4). The timing between birth and final positioning of HCs varied broadly spanning between 9 h and almost 16 h (Figure 3B). Also timing between birth, retraction of the apical process and final position varies (Supplementary Figure 3B). Single cell trajectory analysis revealed that, in contrast to the highly stereotypic migration behaviour of RGCs (Icha et al., 2016), each HC followed its own individual path and migrations tracks took varying amounts of time (Figure 3C). Trajectories are faster and more directed at early migration stages as verified by mean square displacement (MSD) and directionality analysis (Figure 3C and 3D, Supplementary Figure 3C), most likely due to the maintenance of HC apical attachment (Chow et al., 2015). However, trajectories become more diverse and less directed after this stage (Figure 3C and 3D, Supplementary Figure 3C). Tangential migration differed significantly among HCs showing some cells more constrained in nasal-temporal direction and others moving substantially along this axis (Figure 3E). Despite these single cell differences, all HCs robustly reach their correct layer where they start polarizing (Figure 3A and Movie 4).

**Figure 3:**
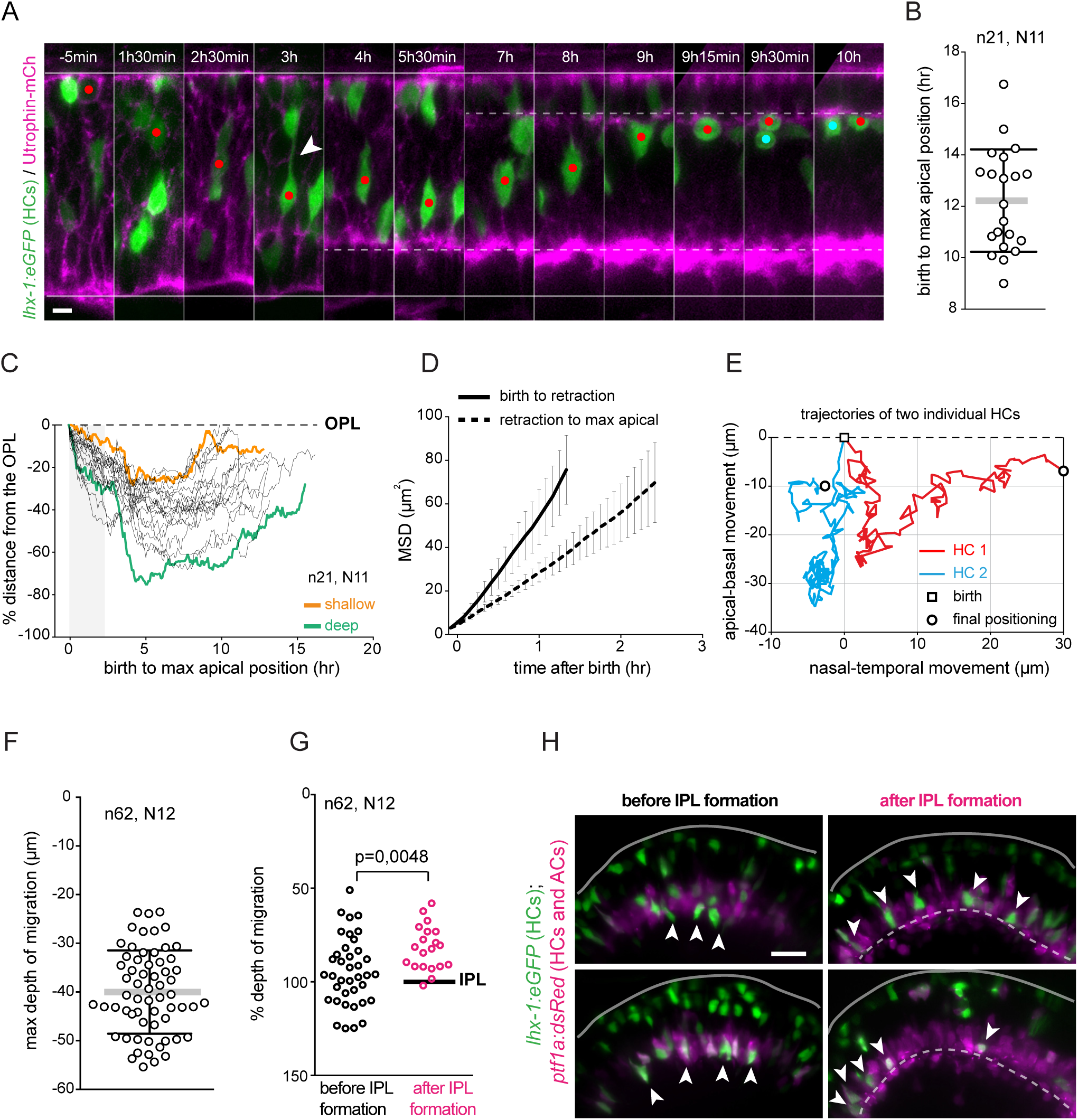
Bidirectional HCpr migration is heterogenous but HCprs always stopover within the amacrine cell layer and heterogeneity in HC depth depends on IPL formation. **(A)** Montage of HC migration from HCpr birth to final positioning. *lhx-1:eGFP* (HCpr and PR, green); *Utrophin-mCherry* (cell outline, magenta), red dot: tracked cell, white arrowhead: apical process before detachment, red and cyan dots sister HCs after terminal division. Time is relative to HC birth. White line: apical surface (top), basal membrane (bottom). White dotted line: OPL (top) and IPL (bottom). Scale bar, 5 µm. **(B)** Scatter plot of HC birth to max apical position. Lines: mean with SD. **(C)** Single cell trajectories of HCprs bidirectional migration from birth to final positioning as % of the total thickness of the retina. orange: example of shallow track, green: example of deep track. Shadow: Initial stage of migration. **(D)** MSDs of HCprs for different phases; from birth to apical process retraction (black) from retraction to final positioning (black dotted). SDs shown. **(E)** Representative examples of HCpr migration along the apical-basal and nasal-temporal retinal axis. **(F)** Distribution of HCpr maximum migration depth. Lines: mean with SD. **(G)** % of HCpr maximum migration depth relative to the retinal thickness, before (black) and after (magenta) IPL (black line) formation. **(H)** Examples of HCpr migration before and after IPL formation. *lhx-1:eGFP* (HCpr and PR, green), *ptf1a:dsRed (*AC and HCpr, magenta). White line: apical surface, dashed white line: IPL. White arrowheads: HCpr position. Scale bar, 10 µm.

### HCs always migrate into the AC layer before apical turning and depth of this migration depends on the presence of the IPL

Similar to other parameters, depth of migration also varied strongly among individual HCs (Figure 3F). To find tissue wide parameters responsible for these differences, we analyzed migration depth in correlation with the formation of the IPL that starts around 54 hpf and contains processes connecting amacrine cells (ACs), BCs and RGCs (Godinho et al., 2005). When defining the distance between the final OPL and IPL as 100% of the migratory path, shallow cells reached less than 50% of this distance, while HCs moving more deeply could surpass the prospective IPL position by 20% (Figure 3F and 3G). However, HC migration deeper than the prospective IPL occurred only before this structure emerged (Figure 3G and 3H). Early HCs that pass the depth of the prospective IPL were nevertheless able to return to apical positions (Figure 3H). Once the IPL was formed however, HCs did not migrate below it (Figure 3G and 3H). Interestingly, regardless of migration depth, all HCs were in contact with the AC layer (Figure 3H and Movie 1) suggesting that migration into the AC layer is necessary for returning apically. As HCs cannot go deeper than the IPL after its formation, this structure most likely acts as a steric hindrance for HCs.

### HC sister cell dispersal varies in time and final nasal-temporal cell distance

Despite the variations found for HC cell cycle and migration parameters, HC layer formation occurred robustly in all embryos imaged (N=21). We thus speculated that sister cell dispersal is important to fill the HC layer independently of single cell behaviour. We expected that this dispersal showed similar heterogeneity as other parameters to buffer the previous stochasticity observed. This was indeed the case (n=20, N=6). While some sister HCs moved substantially in apicobasal direction after terminal divisions and ended up in close nasal-temporal vicinity (Figure 4A and 4B and Movie 5), others moved very little apico-basally but dispersed much more laterally with final positions of sister cell bodies tenth of micrometer apart (Figure 4C and 4D). Lateral HC sister cell movement took between 1h 40 min and 6hr 30 min (Figure 4E) and lateral dispersion differed substantially among cohorts of sister cells upon final mitosis, ranging from 0,6 µm to 29 µm (Figure 4F).

**Figure 4:**
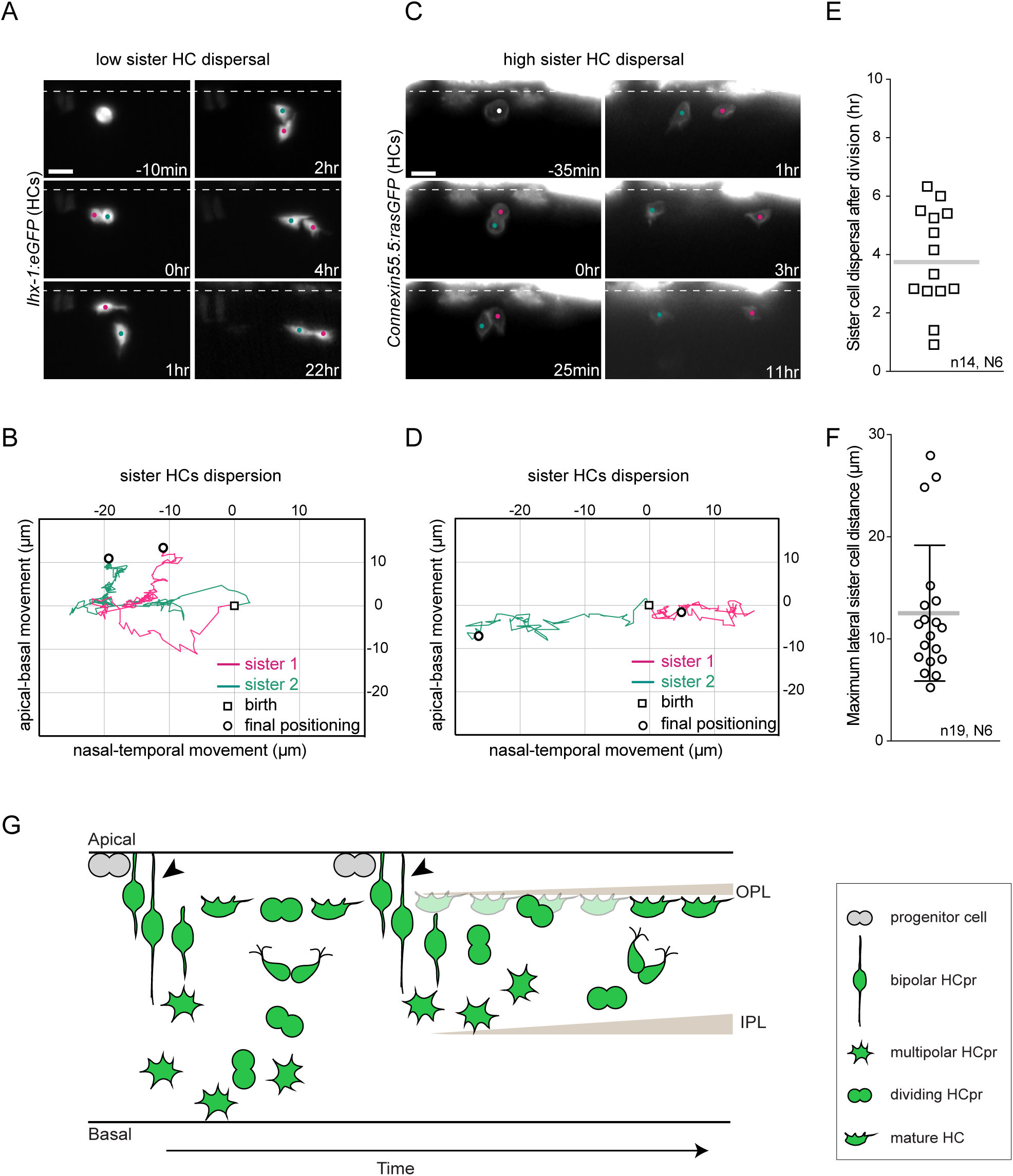
Sister cell dispersal is heterogenous in length and distance between cells. **(A)** Example of sister cell dispersal, *lhx-1:eGFP*, Scale bar 10 µm. **(B)** Trajectories of **(A)**. **(C)** Example of sister cell dispersal, *Connexin:rasGFP*, Scale bar 10µm. **(D)** Trajectories of **(C)**. **(E)** duration of HC sister cell dispersion. Line: mean. **(F)** Maximum lateral distance between sister cells upon final HC mitosis. Line: mean with SD. **(G)** Scheme of HC mitotic and migration behaviour. Retinal progenitors (grey) generate HCprs (green) apically. HCprs first migrate basally while keeping their apical attachment (black arrowheads). Upon losing this attachment, HCprs undergo multipolar migration reaching the basal INL. Depth of basal migration depends on IPL presence (no IPL: deep, IPL: shallow). HCprs invert movement and freely migrate apically to form the HC layer. En route, HCprs undergo a terminal mitosis along the INL.

Thus, like other parameters, sister cell dispersion within the HC layer is highly heterogenous. Interestingly, sister HCs can end up far apart from each other arguing against the idea that functional retinal columns exist, at least when it comes to HC clones (He et al., 2012) as suggested previously for mouse horizontal cells (Reese et al., 1995).

## DISCUSSION

Overall, the use of long-term quantitative light sheet microscopy enabled us to present first evidence that in addition to stochasticity in retinal fate decisions (Baye and Link, 2007; Del Bene et al., 2008; He et al., 2012) also later stages of retinal development, in particular retinal lamination, involve stochasticity. The example presented here, the formation of the HC layer, demonstrates this to an extreme extent: All parameters analyzed over hundreds of cells from many embryos showed heterogeneity. The term stochasticity applies, as single cell behaviour cannot be predicted. Instead, some degree of random chance, seems to govern the cells’ migratory path, division timing and position. Our findings are summarized in Figure 4G.

However, for other cell types that laminate in the retina this is not the case. RGCs laminate via somal translocation and their timing and migration trajectories are highly stereotypic (Icha et al., 2016). What could be the reason for these differences? We speculate that one reason could be the presence or absence of an attached apical process. RGCs are apically attached until they reach the RGC layer and only upon apical process detachment undergo random walk like behaviour during fine positioning (Icha et al., 2016). HCs instead lose their apical attachment early during migration (Chow et al., 2015) (Figure 4G) and from then on navigate freely through the tissue without any predisposed track information. Thus, timing and depth of migration can vary and this heterogeneity needs to be buffered by the heterogeneity of other parameters like sister cell dispersal. This in turn ensures robust HC layer formation. Consequently, while the behaviour of single cells is not predictable, at the tissue level it still produces a robust pattern. The obvious next steps is to understand how this emergence of a well-organized tissue from seemingly chaotic single cell behaviour is ensured. To this end we need to extract the tissue wide parameters guiding it. Such parameters could be coded in chemical guidance cues; however, also mechanical tissue properties could play a role. Follow up studies will therefore open an exciting new array of questions asking which additional neuronal pattern formation phenomena involve stochasticity but also how this single cell stochasticity nevertheless forms reproducible lamination at the tissue level.

## MATERIALS AND METHODS

### Horizontal cell labelling

HCs have been identified in all vertebrate retinas from fish to humans and are divided into two main types; axon-bearing and axon-less. In the adult zebrafish retina, three types of axon-bearing HCs (H1, H2 and H3) and one type of axon-less HC (HB) have been identified (Song et al., 2008). HCs can be labelled by different proteins and transcription factors (TFs) that are often expressed in a subtype-specific manner (Supplementary Figure 1A). In zebrafish, the gap junction proteins Connexin55.5 is expressed in axon-bearing HC subtypes (Godinho et al., 2007; Shields et al., 2007). The pancreas specific transcription factor 1a (Ptf1a) is expressed in all HC subtypes in the zebrafish retina (Jusuf et al., 2011), while the TF *thyroid hormone receptor* β*2* (*tr*β*2*) labels 70% of Ptf1a+ HCs in the zebrafish retina (Suzuki et al., 2013). In the chick retina, the Lim family TFs Islet1 (Isl1) and Lim1/Lhx1 are expressed in axon-less and axon-bearing HC subtypes, respectively (Edqvist et al., 2008). Whether Lim1/Lhx1 has a similar heterogenous expression pattern in HC subtypes in zebrafish is not currently known but we see that a majority but not all Ptf1a positive HCs are labelled by this marker. We here use a combination of these HC markers (Supplementary Figure 1B and C) and show that no marker/subtype-specific HC behaviour is observed (Supplementary Figure 1E).

### Zebrafish work

Zebrafish were maintained and bred at 26.5°C. Zebrafish embryos were raised at 28.5°C or 32°C and staged in hour post fertilization (hpf) according to Kimmel (KIMMEL et al., 1995). Embryos were kept in E3 medium which was supplemented by 0.003% 1-phenyl-2-thiourea (PTU) from 8 ± 1 hpf onwards and renewed daily. PTU was added to prevent pigmentation. Embryos were anesthetized in 0.04% tricane methanesulfonate (MS-222; Sigma-Aldrich) prior to sorting for fluorescent signal (>35 hpf). Refer to S1 Table for a list of transgene lines used in this study. See S3 Table for a list of experiments and number of cells and embryos. All animal work was performed in accordance with the European Union (EU) directive 2010/63/EU as well as the German Animal Welfare act.

### DNA injections

To mosaically label HCs, DNA constructs were injected into the cytoplasm of one-cell stage embryos. Constructs were diluted in double-distilled H_2_O supplemented with 0.05% phenol red (to visualize the injection material) (Sigma-Aldrich). Injected volumes ranged from 1 to 1.5 nl. DNA concentrations were 20-30 ng/µl and did not exceed 30 ng/µl even when multiple constructs were injected. See S2 Table for a list of injected constructs.

### Heat shock of embryos

To induce expression of the heat shock promoter (hsp70) driven constructs and transgenes, embryos were incubated in a water bath 1.5-3 hr before imaging for 20 min at 39°C. See S1 and S2 Tables for a list of heat-shock induced transgenes and constructs, respectively.

### Blastomere transplantations

Transplantation dishes were prepared by floating a plastic template in a Petri dish that was half-filled with 1% low-melting-point agarose in E3. When the agarose solidified, plastic templates were gently removed, leaving an agar mold that contains rows of wells to hold embryos. Embryos at stages high to sphere were dechorionated in pronase (Roche) and dissolved in Danieu’s buffer. Dechorionated embryos were transferred to wells in agarose molds using a wide-bore fire-polished glass pipet. At 1000-cell-stage, cells from the donor embryos were transplanted into the animal-pole of the acceptor embryos using a Hamilton syringe. The animal-pole region is fated to become retina and forebrain tissue. Transplanted embryos were kept on agarose for about 3–5 h and then transferred onto glass dishes, that contained E3 medium supplemented with 0.003% PTU and antibiotics (100 U of penicillin and streptomycin, Thermo Fisher Scientific). Transplanted embryos were identified via fluorescence and imaged from 42hpf (see S3 Table).

### *In vivo* light sheet fluorescent imaging (LSFM)

Imaging started between 35hpf and 48hpf developmental (see S3 Table). Embryos were manually dechorionated and mounted in glass capillaries in 1% low-melting-point agarose as previously described (Icha et al., 2016). The sample chamber was filled with E3 medium containing 0.01% MS-222 (Sigma) (to immobilize embryos) and 0.2 mM PTU (Sigma). Imaging was performed on a Zeiss Light sheet Z.1 microscope (Carl Zeiss Microscopy) with a Zeiss Plan-Apochromat 20× water-dipping objective (NA 1.0) at 28.5°C. Z-stacks spanning the entire eye (100-120 μm) were recorded with 1 μm optical sectioning. Z-stacks were recorded every 5min for 6-42 hours, with double sided illumination mode. The system was operated by the ZEN 2014 software (black edition).

### Image processing and analysis

The raw LSFM data were deconvolved in ZEN 2014 software (black edition, release version 9.0) using the Nearest Neighbor algorithm. Minimal image pre-processing was implemented prior to image analysis, using open source ImageJ/Fiji software (http;//fiji.sc). Processing consisted of extracting image subsets or maximum intensity projections of a few slices. Processed files were then analyzed in Fiji and the data was plotted using MATLAB or Microsoft Excel.

#### 1. Mitosis and cell cycle length analysis

##### Mitosis length analysis

HC rounding was defined as the onset of mitosis and the point of anaphase as the end of mitosis. The time required for this was defined as the mitosis length (Figure 1C and F).

##### Metaphase-to-Anaphase length

To visualize HC nuclei, *Tg(lhx-1:eGFP)* animals were crossed with *Tg(hsp70:H2B-RFP)* animals. Metaphase phase was detected by the condensed and elongated nature of the chromatin in the centre of the cell. Anaphase was assessed by chromatin separation (Supplementary Figure 2C).

##### HC cell cycle period

To assess cell cycle phase of HCs, the Hsp70::EGFP-PCNA DNA plasmid was injected into one-cell stage embryos. Individual PCNA+ HCs were manually followed in Fiji from birth to terminal mitosis. Onset of G1 was defined as the time of HC birth at the apical side. End of G1 was defined as last timepoint prior to PCNA foci appearance (Figure 2A). S phase was defined between the appearance and disappearance of PCNA foci (Figure 2A). G2 was measured from the disappearance of PCNA foci (end of S phase) to cell rounding (Leung et al., 2011).

#### 2. Division Orientation

Division orientation was analysed manually using the angle tool in Fiji RRID:SCR_002285 (Schindelin et al., 2012) from maximum intensity projections of a 5 slices of LSFM images. For each observed anaphase cell, a line was drawn parallel to the division plane, and the angle was measured with respect to the temporal-nasal tissue axis (horizontal image axis) (Supplementary Figure 1D). Data was plotted as a polar histogram in MATLAB.

#### 3. Mitotic position

The Fiji Line tool was used to manually measure the distance from the center of the mitotic HCpr (at rounding) and the OPL in the maximum projection of 3-5 consecutive Z slices (Supplementary Figure 1D).

#### 4. Sample drift correction

First, maximum projected sub-stacks (5 Z slices) of the raw live images were generated in Fiji. XY-drift of 2D stacks was then corrected using a manual drift correction Fiji plugin (http://imagej.net/Manual_drift_correction_plugin, ImageJ).

#### 5. Analysis of HC bidirectional migration, directionality ratio, and MSD analysis

##### Labelling migrating HCs

Three approaches were used to label HCs. 1) double transgenic animals were generated by crossing transgene lines in Table S1: *Tg(lhx-1:eGFP);Tg(ath5:gap-RFP),Tg(lhx-1:eGFP);Tg(*actb2:*mCherry-Hsa.UTRN) and Tg(lhx-1:eGFP); Tg(ptf1a:DsRed*) (Table S3). 2) blastomere transplantation was performed to mosaically label HCs. 3) microinjection of plasmid DNA encoding *trβ2:tdTomato* or *Connexin:55.5:ras*GFP was used to mosaically label HCs in the retina (Table S2 and S3).

##### Tracking migrating HCs

The migrating HCs were manually tracked by following the centre of the cell body in 2D images using MTrackJ plugin in Fiji (Meijering et al., 2012). The resulting trajectories were analysed as described previously (Icha et al., 2016). MSDs and directionality ratios were calculated in the DiPer program (Gorelik and Gautreau, 2014) and executed as a macro in Excel (Microsoft).

#### 6. Maximum depth of HC migration analysis

The relative percentage of maximum HC depth was defined by the apex of HC basal position using individual HC trajectories. At this point, distance of the center of the cell body from the OPL was divided by the OPL to IPL thickness (Supplementary Figure 1F).

### Statistical analysis

All statistical analysis was performed using the Prism software (version 6.0c for Mac OS; GraphPad Software) to create graphs. In total, data were collected from 432 HCs in 24 independent time-lapse imaging experiments (S3 Table). Pearson correlation coefficient was used to examine associations between length, position and angle of HCpr mitosis as well as HCpr cell cycle length and position. The correlation coefficient, r, ranges from −1 to +1, where 1 shows perfect correlation, 0-1 The two variables tend to increase or decrease together, 0 the two variables do not vary together at all, −1-0 one variable increases as the other decreases, −1 perfect negative or inverse correlation.

## ACKNOWLEDGEMENTS

We thank M. Matejcic, I. Yanakieva and the Norden lab for useful project discussions and helpful comments on the manuscript. C. Zechner is thanked for discussion on stochasticity. We are grateful to I. A. Deniz, H. Hollak and S. Kaufmann, the Light Microscopy Facility, the Scientific Computing Facility and the Fish Facility of the MPI-CBG for experimental help. S. Maddu is thanked for help with analysis. We thank R. Wong and L. Godinho for generously sharing constructs.

The authors declare no competing financial interests.

## Supplementary information

### SUPPLEMENTARY MOVIE LEGENDS

**Movie 1: HC lamination during zebrafish retina development.** Light-sheet time-lapse imaging of HC layer formation during retina development. *lhx-1:eGFP* (green) labels HCs and *ptf1a:dsRed* labels both HCs and ACs (magenta). HCs are born at the apical side of the retina and then migrate to the basal INL. After stop-over within the AC layer (magenta), HCs undergo apical migration to form the thin HC layer below the OPL at the apical side of the retina. Imaging started around 42hpf and ended at 70hpf. Time interval 5min. Time in h:min and scale bars = 50 µm. Related to Figure 1 and 3.

**Movie 2: HC exhibit spindle rocking during division.** Time-lapse imaging of HC division dynamics using Tg(*lhx-1:eGFP*); Tg(hsp70:*h2b:RFP). lhx-1:eGFP* (green) labels HCs and H2B-RFP labels nuclei (magenta). Arrow points to the tracked HC. Time interval 5min. Time in h:min and scale bars = 20 µm. Related to Figure 1 and Supplementary Figure 2.

**Movie 3: HC cell cycle in a developing retina.** Light-sheet time-lapse imaging of PCNA dynamics from HC birth to its final cell cycle. Mosaic expression of *hsp70:GFP-PCNA* (magenta, nuclei) and *Trβ2:tdTomato* (green, HCs and PRs). White arrowhead points to the tracked HC. Cyan and red arrowheads point to HC sister cells after HC final division. Time interval 5min. Time in h:min and scale bars = 10 µm. Related to Figure 2.

**Movie 4: HCs undergo a bidirectional migration.** Light-sheet time-lapse imaging of HC migration dynamics from birth to final positioning. Mosaic expression of *connexin55.5:rasGFP* (green, HCs) and *hsp70:H2B-RFP* (magenta, nuclei). White arrowhead points to the tracked HC. Cyan and red arrowheads follow HC sister cells from division to final positioning. Time interval 5min. Time in h:min and scale bars = 10 µm. Related to Figure 3 and Supplementary Figure 3A-C.

**Movie 5: HC sister cells undergo heterogenous dispersion.** Light-sheet time-lapse imaging of HC sister cells from birth (HC final division) to final positioning. Transplanted *lhx-1:eGF* (grey, HCs and PRs) into a control retina (left) shows that two sister cells end up at close vicinity to each other. Mosaic expression of *connexin55.5:rasGFP* (grey, HCs) shows that two sister cells final position is far from each other. White arrowhead points to the tracked HC before its division. Cyan and red arrowheads follow HC daughter cells from division to final positioning. Time interval 5min. Time in h:min and scale bars = 10 µm. Related to Figure 4 and Supplementary Figure 3D.

## SUPPLEMENTARY FIGURE LEGENDS

**Supplementary Figure 1:**
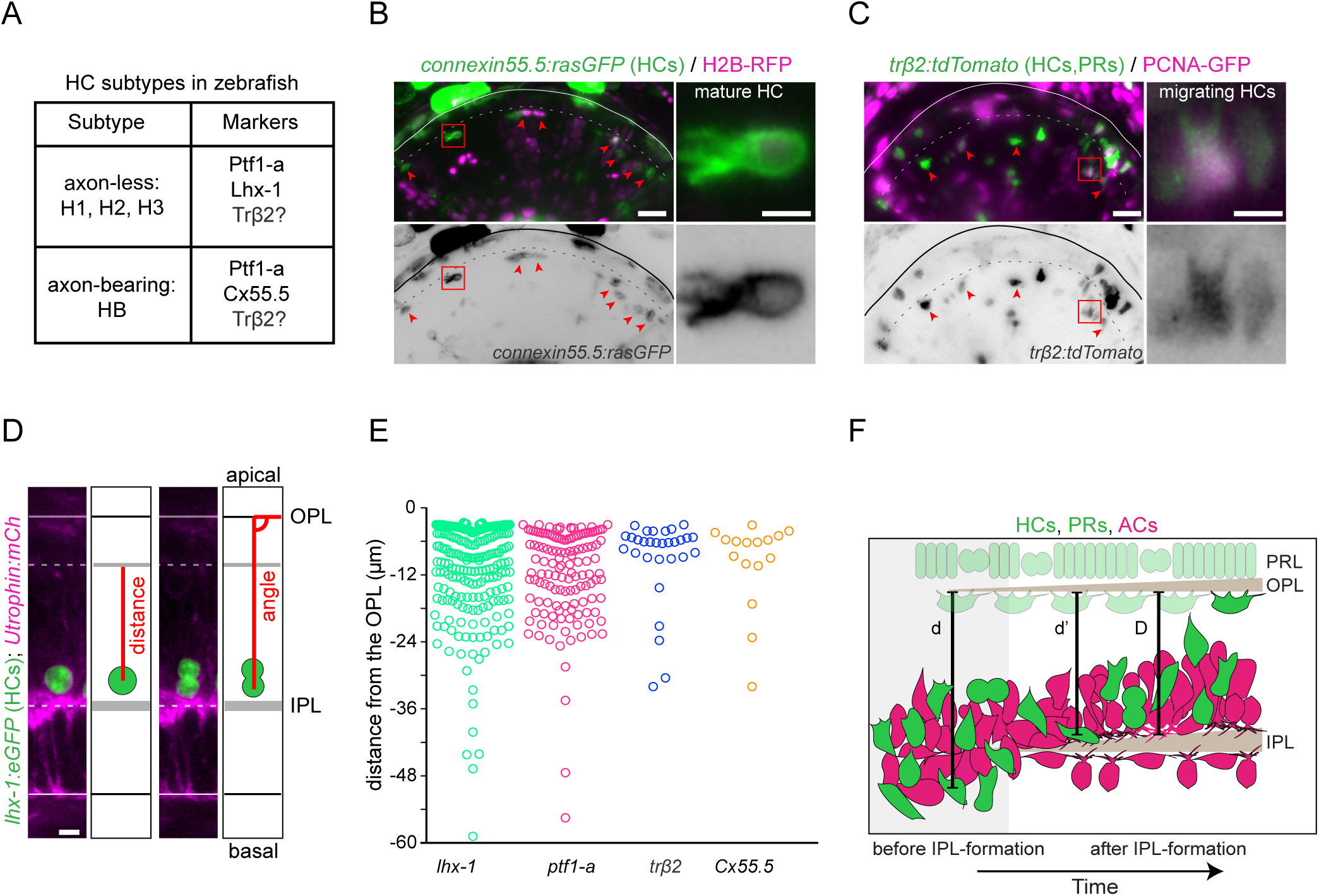
**(A)** HCs subtypes and markers in the zebrafish retina. **B-C)** Typical examples of HCprs labelled with **(B)** *Connexin55.5:rasGFP* (Cx55.5) and **(C)** *Trβ2:tdTomato*. Red arrowheads point to HCs within the HC layer. Dashed line represents the OPL. Scale bar 10µm. Insets of the red-boxed region shows magnified HC. Scale bar 5µm. **(D)** Schematic showing measurement of HC distance from the OPL at rounding (left) and HC angle of division (right). Scale Bar, 5 µm. **(E)** Position of mitotic HCprs for different HC marker populations. **(F)** Schematic showing depth of migration measurements. Relative thickness of the INL was quantified by drawing a line (D) from the OPL to the IPL using Fiji Line tool. To quantify HC position, a line (d’) was drawn from the center of HC to the OPL. Relative percentage of HC depth was defined by dividing d’/D. For earlier developmental stages before the IPL formation, thickness of the INL was measured once the IPL was formed within the same region.

**Supplementary Figure 2:**
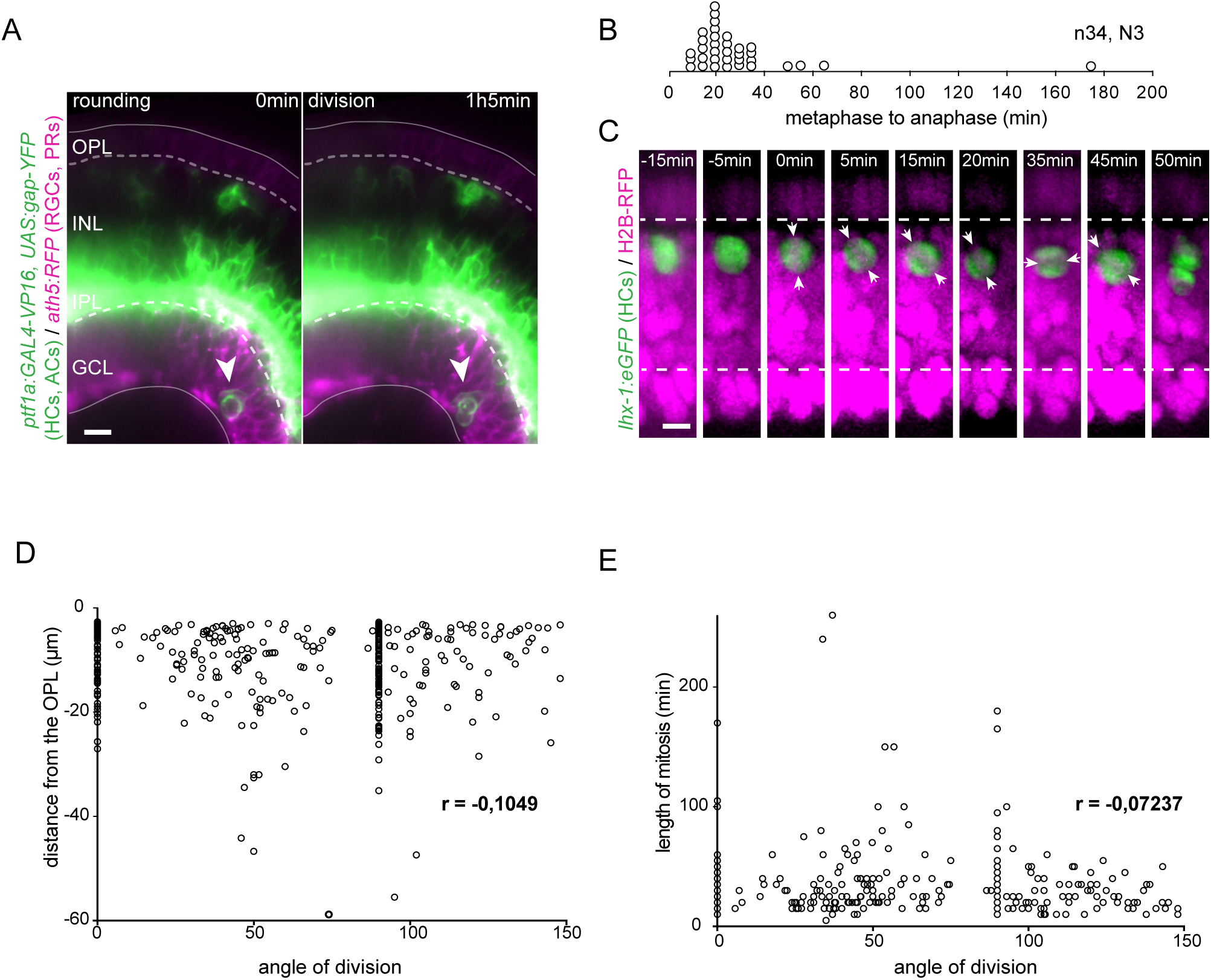
Heterogeneity in mitotic position and behaviour of committed HCpr subtypes. **(A)** Rare example of HCpr dividing in the RGC layer. *ptf1a:GAL4-VP16, UAS:gap-YFP* (green, HCpr and AC); *ath5:RFP (PR and RGC).* White arrow: mitotic HCpr. White dashed line: OPL (top), IPL (bottom). Scale Bar, 10 µm. **(B)** Spread of metaphase to anaphase duration in mitotic HCprs. **(C)** Example of HCpr spindle rocking before mitosis. *lhx-1:eGFP* (HCpr, green); *H2B-mcherry (nuclei, magenta), white arrows: position of metaphase plate.* White dashed line: the OPL (top), the IPL (bottom). Scale Bar, 5 µm. **(D)** Distribution of HCpr mitotic angle versus position does not show a strong correlation. **(E)** Distribution of HCpr angle versus length of mitosis does not show a strong correlation.

**Supplementary Figure 3:**
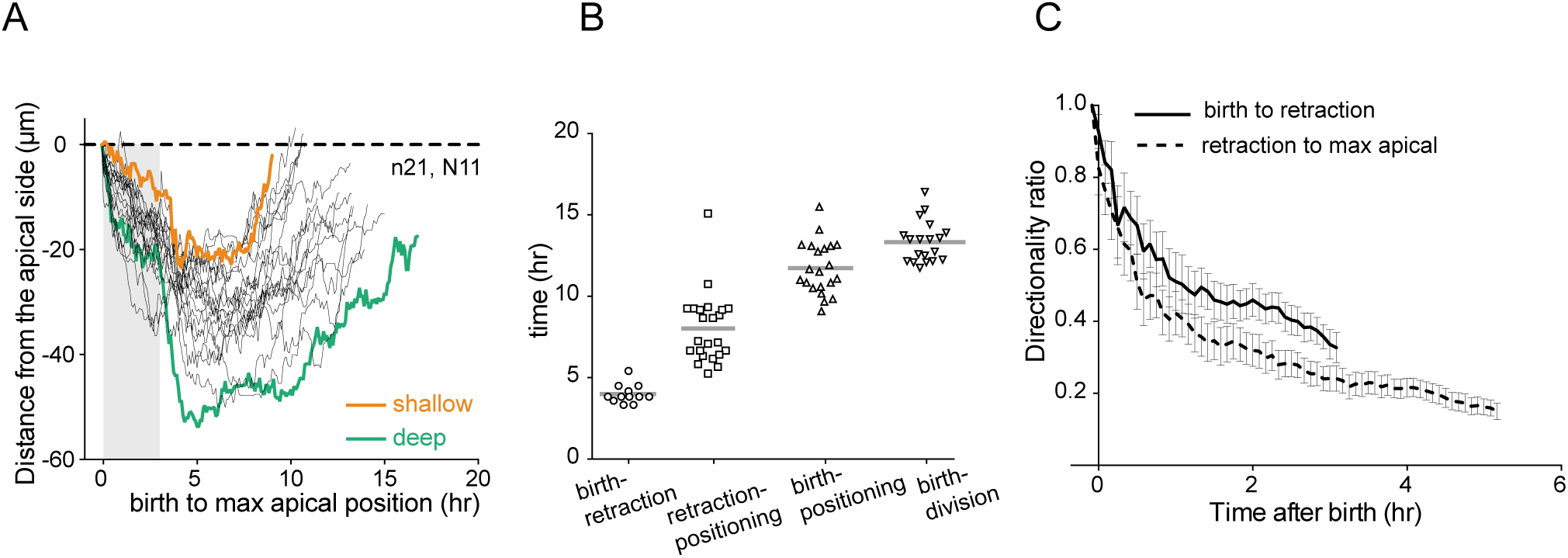
HCprs migration is heterogenous. **(A)** Single cell trajectories of HCs from birth to final positioning. This graph depicts distance of migrating HCs shown in Figure 3C in micron from apical side. **(B)** Scatter plot showing length of different phases of HC migration. **(C)** Directionality plots of HCs for different phases of migration; from birth to retraction of the apical attachment (black line, first 1hr33min) and from retraction to final positioning (black dotted line, first 2hr40min).

## SUPPLEMENTARY TABLES

**Table S1:** Zebrafish lines corresponding to images and plots in figures

| Line | Structures labeled | Figure | Reference |
| --- | --- | --- | --- |
| *Tg(SoFa2)* | Membranes of all retinal cells | 1C, E-J<br>Suppl. 2A, D, E | (Almeida et al., 2014) |
| *Tg(ptf1a:Gal4-VP16, UAS:gap-YFP)* | Membranes of ACs and HCs | 1C, E-J<br>Suppl. 2D-E | (Pisharath and Parsons, 2009)<br>(Williams et al., 2010) |
| *Tg(ptf1a:DsRed)* | ACs and HCs | 1B, D, D'<br>3H | (Vitorino et al., 2009) |
| *Tg(lhx-1:eGFP)* | HCs and PRs | 1B, D-J<br>3A-H<br>4A, E-F<br>Suppl. 2C<br>Suppl. 3 | (Swanhart et al., 2010)<br>(Boije et al., 2015) |
| *Tg(hsp70:H2B-RFP)* | Chromatin | Suppl. 1B<br>Suppl. 2B-C | (Dzafic et al., 2015) |
| *Tg(βactin:maKate2-ras)* | All membranes | 1E-J<br>3B-D | (Matejčić et al., 2018) |
| *Tg(ath5:gap-RFP)* | Membranes of RGCs and PRs | 1B<br>Suppl. 2A | (Zolessi et al., 2006)<br>(Icha et al., 2016) |
| *Tg(actb2:mCherry-Hsa.UTRN)* | F-Actin | 3A | (Compagnon et al., 2014) |

**Table S2:** List of constructs corresponding to images and plots in figures

| Construct | Structures labeled | Figure | Reference |
| --- | --- | --- | --- |
| *trβ2:tdTomato* | HCs and PRs | 2A-E<br>Suppl. 1C, E | (Suzuki et al., 2013) |
| *hsp70:PCNA-GFP* | Cell cycle phase marker | 2A-E<br>Suppl. 2C | (Icha et al., 2016) |
| *Connexin:55.5:rasGFP* | Membrane of HCs | 4C, E,D,F<br>Suppl. 1B | (Weber et al., 2014) |
| *hsp70:H2B-RFP* | Chromatin | Suppl. 1B | (Strzyz et al., 2015) |

**Table S3:** List of *in vivo* time lapses used in this study

| Movie | Transgenes/ constructs injected | analyzed HC | Time-lapse started at (hpf) | Duration of time-lapse (hr) |
| --- | --- | --- | --- | --- |
| 1 | *Tg(SoFa2)* | 24 | 36 | 40 |
| 2 | *Tg(SoFa2)* | 18 | 42 | 34 |
| 3 | *Tg(SoFa2)* | 48 | 40 | 38 |
| 4 | *Tg(ptf1a:Gal4-VP16, UAS:gap-YFP)* | 32 | 35 | 42 |
| 5 | *Tg(ptf1a:Gal4-VP16, UAS:gap-YFP)* | 11 | 42 | 35 |
| 6 | *Tg(ptf1a:Gal4-VP16, UAS:gap-YFP)* | 3 | 48 | 10 |
| 7 | *Tg(lhx-1:eGFP); Tg(ptf1a:DsRed)* | 86 | 40 | 40 |
| 8 | *Tg(lhx-1:eGFP)* transplanted into WT | 12 | 42 | 40 |
| 9 | *Tg(lhx-1:eGFP)* transplanted into *Tg(βactin:maKate2-ras)* | 8 | 42 | 40 |
| 10 | *Tg(lhx-1:eGFP); Tg(ath5:gap-RFP)* | 37 | 42 | 35 |
| 11 | *Tg(lhx-1:eGFP); Tg(ath5:gap-RFP)* | 6 | 48 | 24 |
| 12 | *Tg(lhx-1:eGFP); Tg(actb2:mCherry-Hsa.UTRN)* | 31 | 48 | 30 |
| 13 | *Tg(lhx-1:eGFP); Tg(actb2:mCherry-Hsa.UTRN)* | 31 | 48 | 35 |
| 14 | *Tg(lhx-1:eGFP); Tg(hsp70:H2B-RFP)* | 33 | 42 | 30 |
| 15 | *Tg(lhx-1:eGFP); Tg(hsp70:H2B-RFP)* | 4 | 42 | 31 |
| 16 | *trβ2:tdTomato* and *hsp70:PCNA-GFP* were co-injected. | 1 | 42 | 11 |
| 17 | *trβ2:tdTomato* and *hsp70:PCNA-GFP* were co-injected. | 11 | 48 | 20 |
| 18 | *trβ2:tdTomato* and *hsp70:PCNA-GFP* were co-injected. | 4 | 42 | 20 |
| 19 | *trβ2:tdTomato* and *hsp70:PCNA-GFP* were co-injected. | 8 | 42 | 40 |
| 20 | *trβ2:tdTomato* and *hsp70:PCNA-GFP* were co-injected. | 5 | 42 | 35 |
| 21 | *trβ2:tdTomato* and *hsp70:PCNA-GFP* were co-injected. | 1 | 48 | 24 |
| 22 | *trβ2:tdTomato* and *hsp70:PCNA-GFP* were co-injected. | 1 | 42 | 28 |
| 23 | *Connexin:55.5:rasGFP* and *hsp70:H2B-RFP* were co-injected. | 8 | 42 | 31 |
| 24 | *Connexin:55.5:rasGFP* and *hsp70:H2B-RFP* were co-injected. | 9 | 42 | 30 |

